# Computationally Guided Design of Dipeptidyl Peptidase-4 Inhibitors

**DOI:** 10.1101/772137

**Authors:** Lauren C. Reynolds, Morgan P. Connolly, Justin B. Siegel

## Abstract

The Type 2 Diabetes Mellitus (T2DM) epidemic undoubtedly creates a need for the development of new pharmaceuticals. With the goal of generating new therapeutics for this disease, computational studies were conducted to design novel dipeptidyl peptidase-4 (DPP-4) inhibitors. Two candidates, generated by chemical intuition-driven design and bioisosteric replacement, were found to have better docking scores than anagliptin, a currently available diabetes medication.

## INTRODUCTION

Type 2 Diabetes Mellitus (T2DM) is dubbed the “epidemic of the 21st century”— affecting millions of patients worldwide. In the United States alone, according to a 2017 U.S. National Health Interview Survey, 23.1 million adults were diagnosed with diabetes and roughly 13.7 million were between the ages of 18-64.^1^ Diabetes was also deemed the 7^th^ leading cause of death in the United States in 2017.^2^ The prevalence of this disease has prompted the pharmaceutical industry to focus on developing therapies for affected patients.^3^ Some currently available T2DM therapies are known to cause side effects such as hypoglycemia and significant weight gain, which serves as the motivation to create improved drug therapies in this area.^3^

Recently, pharmaceutical companies are interested in the drug target dipeptidyl peptidase-4 (DPP-4) since it improves glycemic homeostasis without putting the patient at increased risk of hypoglycemia.^4^ DPP-4 is a serine protease which is responsible for bioactive peptide regulation and the maintenance of glucose levels in the human body.^4^ This enzyme naturally cleaves N-terminal dipeptides from substrates, such as incretin hormones, which are crucial participants in the gut-brain axis.^3^ The inhibition of DPP-4 permits the prolonged availability of incretin hormones such as glucagon-like peptide-1 (GLP-1) and glucose-dependent insulinotropic polypeptide (GIP), allowing for a decrease in the patients’ blood glucose levels. By harnessing this biochemical signaling pathway, the industry has developed approved, DPP-4-targeting drugs such as alogliptin, anagliptin, linagliptin, and sitagliptin.

Though these aforementioned drugs are largely successful in treating the T2DM population, there are various pharmacodynamic and pharmacokinetic properties that are a cause for concern. According to the FDA, alogliptin users are potentially at greater risk for heart failure, especially those with preexisting heart or kidney diseases.^5^ Other side effects of this medication include severe joint pain and pancreatitis.^5^ Pancreatitis and increased risk of pancreatic cancer have also been a topic of debate in recent years for other gliptins as well. A recent meta-analysis seems to suggest a possible increased pancreatitis risk for sitagliptin users.^6^ One study even suggested that pancreatitis was greater than 6-fold more likely to be reported from sitagliptin users versus other T2DM therapies.^7^ Conclusions regarding the long-term effects, such as cancer, are still ongoing since these medications are relatively new. With these adverse effects in mind, the field is still exploring new therapies in an effort to minimize patient risk during treatment.

In an effort to accomplish this task, some computational studies have been conducted to design new DPP-4 inhibitors. These studies use various strategies, such as high-throughput screening or peptidomimetics, to generate inhibitors from a variety of chemical classes.^8^ Despite this, all inhibitors that have activity for inhibiting DPP-4 share several key characteristics. A study using quantitative structure activity relationships (QSAR) revealed that three hydrogen bond acceptors, one hydrogen bond donor, and one aromatic feature are required for activity with DPP-4.^8^ Their model predicted that a bulky group around SER376 in the active site and an electronegative moiety around GLU866 would lead to high activity.^8^ Another study by Kalhotra et al. used a FlexX algorithm to evaluate natural products and their role in DPP-4 inhibition.^9^ They discovered that the natural compound, chrysin, exhibited a molecular docking score of −21.4 kJ/mol and interacted with the catalytic triad site (SER630, ASP708, HIS740) on DPP-4.^9^ Chrysin was evaluated further in vitro, revealing its concentration-dependent inhibition of DPP-4.^9^ Together, these studies show the potential for generating successful alternative therapies.

Here we describe the design of two novel DPP-4 inhibitors and compare them to existing T2DM drugs based on their ADMET (adsorption, distribution, metabolism, excretion, and toxicity) characteristics, chemical properties, and docking scores. Both designed inhibitors exhibited better docking scores than anagliptin while satisfying ADMET criteria, which is a promising starting point for other T2DM mediation alternatives.

## METHODS

Human dipeptidyl peptidase-4 (DPP-4) complexes with corresponding reversible inhibitors were obtained from the RCSB Protein Data Bank website.^10^ Crystal structures of DPP-4 in complex with anagliptin (3WQH)^11^, alogliptin (3G0B)^12^, linagliptin (2RGU)^13^, and sitagliptin (1X70)^14^ were used in subsequent computer modeling software.

PyMOL was used to visualize ligand-protein interactions and measure interaction distances.^15^ ADMET properties were obtained via OpenEye FILTER, which screens for the number of stereocenters, toxicophoric groups, potential for aggregation, bioavailability score, and Lipinski’s rules of 5.^16^ vBrood 2.0 was utilized to build a new drug query via bioisosteric replacement using anagliptin as the lead molecule.^17^

Gaussview and Gaussian 03 allowed for the geometrical creation and semi-empirical optimization of the designed drug candidates.^18^ Conformer libraries of designed compounds were generated using OMEGA.^19^ Drug candidates were then docked into the active site of DPP-4 (PDB ID 3WQH) using FRED and FRED Receptor.^20^ In FRED Receptor, the molecular probe was used to define the active site. For Site Shape Potential, medium quality was selected. The amino acids automatically identified by the program were appropriately labeled as hydrogen bond donors or acceptors under the Protein tab.

## RESULTS AND DISCUSSION

### Evaluation of Known Dipeptidyl Peptidase-4 Inhibitors

An analysis of known dipeptidyl peptidase-4 inhibitors (alogliptin, anagliptin, linagliptin, and sitagliptin) was carried out in order to gain insight on the general characteristics present in successful molecules. It was hypothesized that the improvement of pre-existing molecule’s docking scores or properties could lead to new, innovative inhibitors.

First, each of the inhibitors was docked into the DPP-4 active site using FRED Receptor to generate Chemgauss3 scores. Chemgauss3 provides information on the docking score, steric hindrance, desolvation, hydrogen bond acceptor (HBA), and hydrogen bond donor (HBD) properties (Table 1). Alogliptin was found to have the best docking score at −78.471. Anagliptin’s hydrogen bond donor and acceptor scores were the most prominent at −12.199 and −13.557 respectively. Both linagliptin and sitagliptin showed moderate docking scores; however, their hydrogen bond donor and acceptor capabilities were limited in comparison to alogliptin and anagliptin. Alogliptin and linagliptin had similar steric scores, which is indicative of the number of heavy atoms in the protein that make contact with the drug’s heavy atoms. The Chemgauss3 steric score accounts for van der Waals contacts and protein desolvation energy during protein-ligand binding. The steric score, which accounts for van der Waals contacts and protein desolvation energy during protein-ligand interaction was the least negative in anagliptin though the desolvation score was the highest, which is a penalty the program assigns when the HBD and HBA on the ligand are prevented from interacting with water molecules in the active site.

**Table 1.** Chemgauss3 Scores of Known Drug Molecules Alogliptin, Anagliptin, Linagliptin, and Sitagliptin.

| Name | Docking Score | Steric | Desolvation | Acceptor | Donor |
| --- | --- | --- | --- | --- | --- |
| Alogliptin | -78.471 | -70.995 | 6.983 | -12.170 | -2.289 |
| Anagliptin | -65.076 | -57.903 | 18.582 | -13.557 | -12.199 |
| Linagliptin | -69.651 | -70.249 | 6.950 | -3.073 | -3.279 |
| Sitagliptin | -68.208 | -66.096 | 6.173 | -1.904 | -6.382 |

Another means of evaluating these known inhibitors was through the use of OpenEye FILTER, which analyzes the ADMET properties of the molecules. The ADMET property table generated from the OpenEye command prompt revealed that none of the four drugs violated any of Lipinski’s rules. This, along with the assigned bioavailability score, corroborates the fact that all of these drugs are orally bioavailable. Additionally, all of these molecules have two or fewer chiral centers and no toxicophoric groups, which reduces the probability of adverse effects in the biological system. None of the drug molecules were found to contain toxicophoric groups or exhibit aggregator qualities. In terms of drug discovery, aggregators tend to give false positive results during high-throughput screening (HTS) which can give the impression that these are good lead compounds when in fact they are not (since they may lead to pharmacodynamic and pharmacokinetic issues).^21^

After evaluation of ADMET and Chemgauss3 scores for the known candidates, it was necessary to establish a control for use in further computational studies. Though it would be rational to model the drug candidates after the natural ligands, GLP-1 (MW = 3297.7 g/mol) and GIP (MW = 4983.6 g/mol), these molecules are far too large to be components in an orally-administered drug because of bioavailability constraints. Thus, in order to increase the probability of finding successful drug candidates, anagliptin was chosen because it had the most promise for improvement due to it possessing the highest docking score, having the highest steric score, and lacking the maximal amount of hydrogen bond donors and acceptors (according to Lipinski’s rule).

To visualize and measure the interactions anagliptin makes with the active site, its crystal structure was studied further in PyMOL (Figure 1). Anagliptin (purple) is shown to bind non-covalently to the active site by reacting with nearby amino acids (orange) that make up the protein (green). The strongest interactions made with DPP-4 include: hydrogen bonding with TYR547, GLU 205, GLU206, and pi-pi stacking with PHE357; these bonds are shown with dashed yellow lines and measure 3.3 Å, 3.3 Å, 2.9 Å, and 3.7 Å respectively.

**Figure 1.**
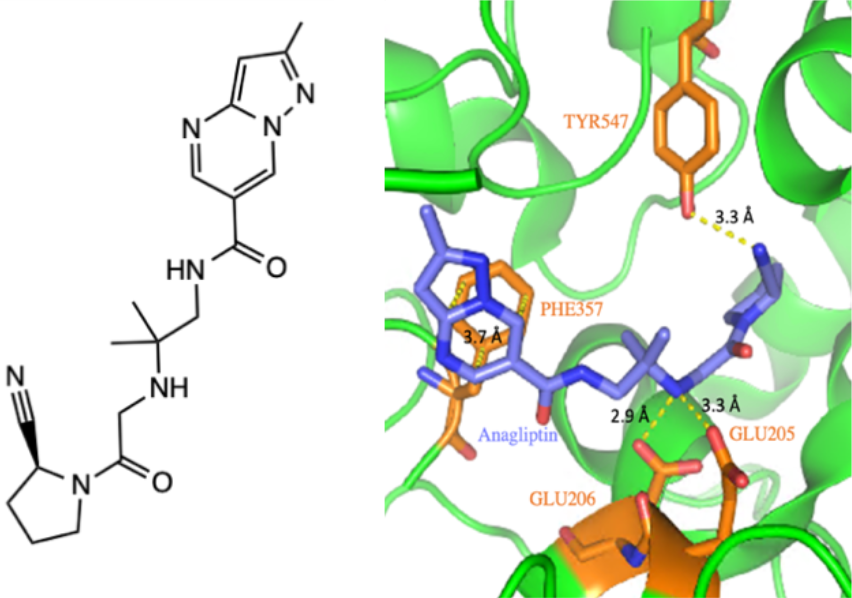
2D Structure of Anagliptin (left) and Anagliptin’s Interactions with TYR547, GLU 205, GLU206, and PHE357 in the DPP-4 Active Site (right).

These crucial interactions for recognition were taken into consideration when selecting sites for alteration via bioisosteric replacement. It was noted that the amide near the fused five and six-membered rings did not have any strong interactions within the active site. Additionally, it was observed that there were limited contacts to the five-membered ring with the cyano functionality. These regions on anagliptin were areas selected for improvement by computational methods.

### Computationally Driven Drug Design

Computational drug design began with loading anagliptin into vBrood 2.0. The five-membered ring with the cyano moiety was altered in an attempt to create new active site interactions and achieve a better overall docking score. After generating a vBrood hit list for this alteration, Favorites, which included **Candidate 1**, were then docked into the DPP-4 active site using FRED Receptor. **Candidate 1** possessed a 2-cyano-3,4-chloro-benzene ring which replaced the preexisting five-membered ring (Figure 2A).

**Figure 2.**
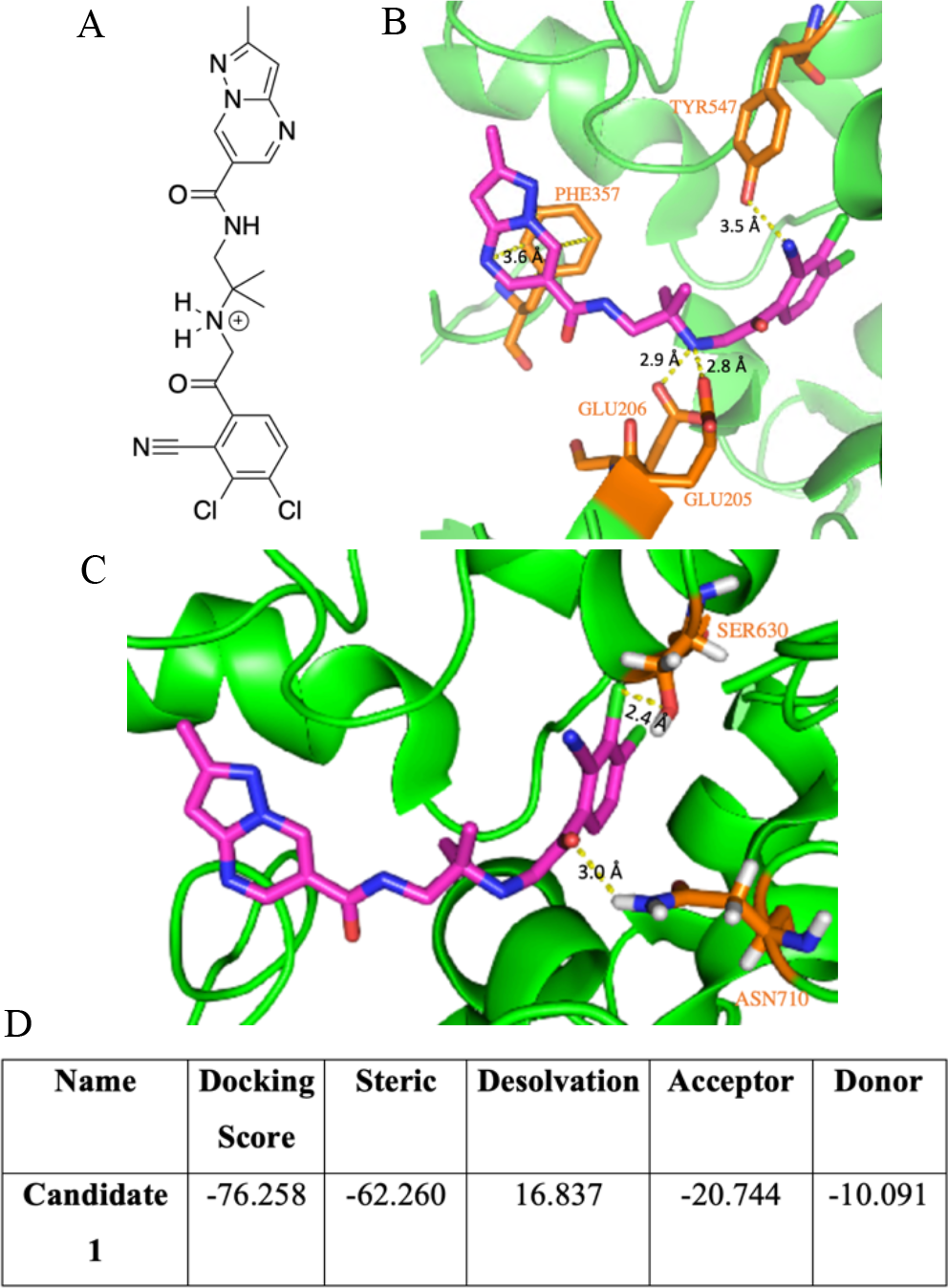
Structure and Interactions of **Candidate 1** with DPP-4 Active Site. A) 2D Structure of **Candidate 1.** B) **Candidate 1**’s Interactions with TYR547, GLU 205, GLU206, and PHE357. C) New Interactions Between **Candidate 1** and SER630 and ASN710 in the Active Site. D) Chemgauss3 Scores of **Candidate 1**.

The docking score obtained for this molecule is −76.258, which is a 11.182 improvement from anagliptin’s docking score (Table 2D). The molecule also had a lower steric score, which is indicative of less unfavorable interactions within the active site. In addition, the desolvation penalty was minimized despite having more acceptor capabilities.

**Table 2.**
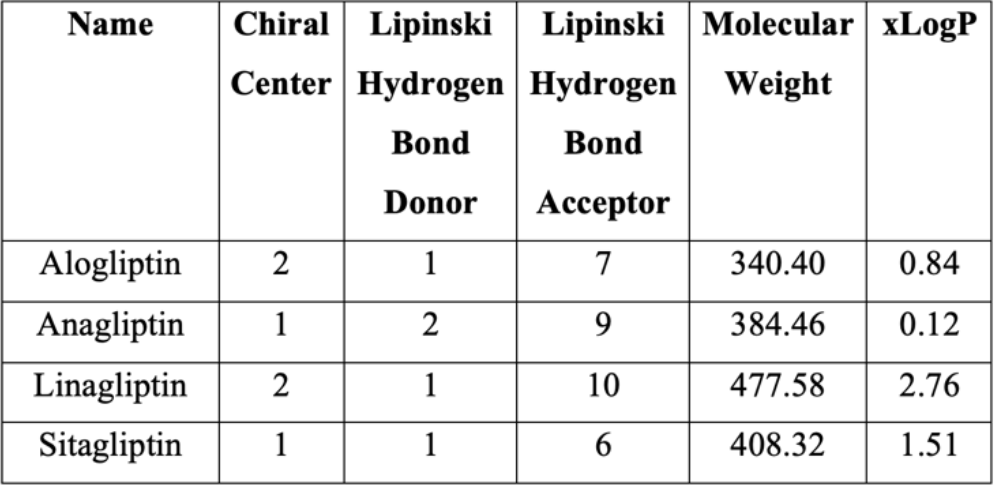
ADMET Properties of Known DDP-4 Inhibitors.

| Name | Chiral Center | Lipinski Hydrogen Bond Donor | Lipinski Hydrogen Bond Acceptor | Molecular Weight | xLogP |
| --- | --- | --- | --- | --- | --- |
| Alogliptin | 2 | 1 | 7 | 340.40 | 0.84 |
| Anagliptin | 1 | 2 | 9 | 384.46 | 0.12 |
| Linagliptin | 2 | 1 | 10 | 477.58 | 2.76 |
| Sitagliptin | 1 | 1 | 6 | 408.32 | 1.51 |

To rationalize why **Candidate 1** had an improved docking score, PyMOL was used to visualize this molecule within the active site. First, the interactions with TYR547, GLU 205, GLU206, and PHE357 were remeasured to evaluate their retention after replacement (Figure 2B). The pi-pi stacking interaction became 0.1 Å shorter. The hydrogen bond between GLU206 maintained its length of 2.9 Å; however, the hydrogen bond with GLU205 became 0.5 Å shorter, which allows for stronger electrostatic interaction. The interaction between TYR547 became 0.2 Å longer, but this can be justified by the fact that the ligand became nearer to the GLU205 residue. Overall, the newly created drug kept the interactions that were originally present on anagliptin.

**Candidate 1** also created new interactions with the active site; these interactions include hydrogen bonding with ASN710 and halogen bonding with SER630 (Figure 2C). The slight downward shift toward the GLU205 and GLU206 residues allowed for the closer proximity of the carbonyl nearest the newly-added benzene ring and ASN710. An additional hydrogen bond in the active site undoubtedly has an impact on docking score improvement. Additionally, the chlorine moiety nearest the cyano group on the ring can participate in halogen bonding with SER630. Halogen bonding is common in pharmaceutical applications since many pharmaceuticals contain halogens.^22^ Halogens, which are elements with high electron-density due to their electronegative behavior, commonly interact with nearby halogen-bond acceptors, which can be O, N, S, or aromatic rings.^22^ Such interactions are used in the pharmaceutical industry to create an improved binding affinity without disturbing other active site interactions.^22^ The addition of the 2-cyano-3,4-chloro-benzene ring allows for a proximal interaction between chlorine and the oxygen on SER630.

Though **Candidate 1** had improved docking scores, it was still very important to evaluate its ADMET properties. Thus, OpenEye FILTER was used to generate its scores (Table 3). Compared to anagliptin, the xLogP increased dramatically, though it is still in the range for oral bioavailability. This increase is due to getting rid of the pyrrolidine ring and replacing it with a benzene; the benzene ring is more nonpolar and hydrophobic. This addition did not violate any of Lipinski’s rules of 5, and it did not illustrate aggregatory behavior. Along with the docking score results, these ADMET properties seem to suggest this drug being a good candidate.

**Table 3.**
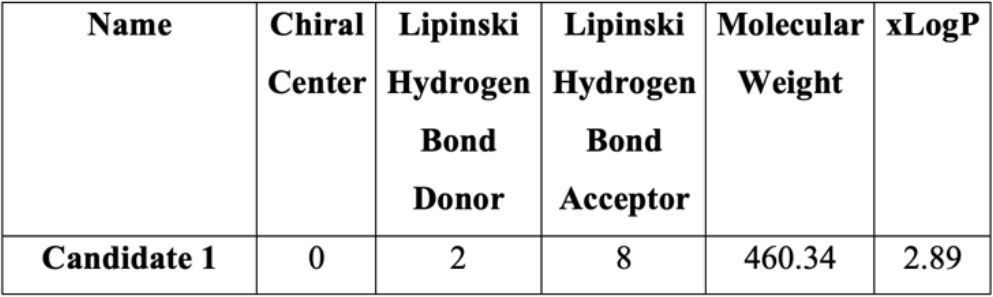
ADMET Properties of **Candidate 1**.

As a whole, vBrood 2.0, FRED Receptor, OpenEye, and PyMOL each aided in the successful creation, modeling, and testing of a novel drug lead **Candidate 1**. Next, a chemical intuition-driven approach will be taken to design one more drug candidate.

### Chemical Intuition Driven Drug Design

During the initial analysis of anagliptin in the active site, it was discovered that the amide by the fused five and six-membered rings did not have any strong interactions. The nitrogen and oxygen in this area could have the potential for hydrogen bonding, but it was clear that the distance of these atoms from appropriate residues was too far. Thus, it was hypothesized that adding a nitrogen ring could increase the steric bulk, potentially allowing for its interaction with a hydrogen bond acceptor or donor. The carbonyl was converted to an alcohol and placed on the five-membered ring to increase its accessibility. It was hypothesized that this modification could undergo hydrogen-bond interactions while will keeping the same initial atoms present. The designed molecule, **Candidate 2**, was built in Gaussview and geometrically optimized in Gaussian (Figure 3A).

**Figure 3.**
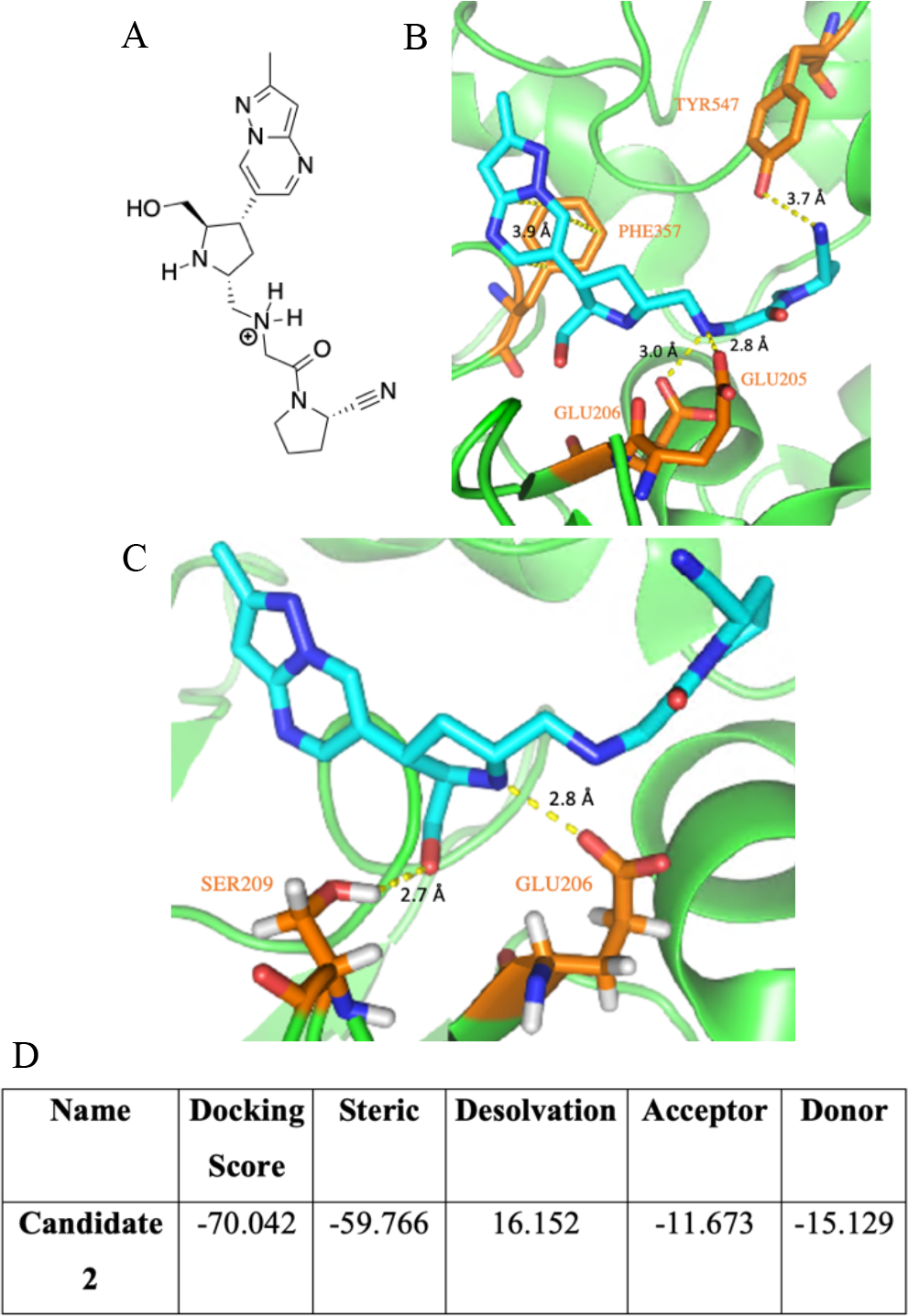
Structure and Interactions of **Candidate 2** with DPP-4 Active Site. A) 2D Structure of **Candidate 2.** B) **Candidate 2**’s Interactions with TYR547, GLU 205, GLU206, and PHE357. C) New Interactions Between **Candidate 2** and SER209 and GLU206 in the Active Site. D) Chemgauss3 Scores of **Candidate 2**.

**Candidate 2** was next docked using the FRED Receptor software, and a Chemgauss3 scoring chart was generated (Figure 3D). A docking score of −70.042 was obtained, which is a 5-point improvement on the docking score of anagliptin. The steric score also improved by roughly 2 points, which could be attributed to the addition of the 5-membered ring allowing it to fill more space within the active site. The donor score also increased by roughly 2 points—probably due to the changing of the carbonyl to an alcohol.

PyMOL was used to determine whether **Candidate 2** maintained the interactions initially present in anagliptin (Figure 3B). The hydrogen bonds to GLU205 and GLU206 stayed relatively unchanged in terms of bond distance. The pi-pi interaction with PHE357 was still present as well—though this length increased by 0.2 Å. The most noteworthy difference was the lengthening of the TYR547-nitrogen interaction by 0.4 Å, which could be too long to be considered a hydrogen bond. Despite this loss of a stabilizing interaction, other new interactions were made, such as a hydrogen bond with SER209 and an additional hydrogen bond with GLU206 to the nitrogen in the ring (Figure 3C). The chemical intuition-driven designed drug undeniably exposes the drug to more hydrogen bond opportunities.

Lastly, the ADMET properties were found via OpenEye FILTER. As expected, the Lipinski hydrogen bond donor value increased by one due to the addition of the alcohol; the hydrogen bond acceptor value remained unchanged as anticipated since the same number of oxygen and nitrogen atoms are present. The xLogP value decreased slightly—possibly due to the increased accessibility of the -OH to water. No toxicophoric groups were present on the molecule, though the presence of four chiral centers could pose potential problems during metabolism. Further testing will need to be conducted to determine the safety of the enantiomers of each stereocenter.

**Table 4.** ADMET Properties of **Candidate 2**.

| Name | Chiral Center | Lipinski Hydrogen Bond Donor | Lipinski Hydrogen Bond Acceptor | Molecular Weight | xLogP |
| --- | --- | --- | --- | --- | --- |
| **Candidate 2** | 4 | 3 | 9 | 398.48 | -0.99 |

Overall, the chemical intuition-driven drug molecule **Candidate 2** seems to show promise since it has an improved docking score due to increased hydrogen-bonding capabilities with SER209 and GLU206.

## CONCLUSION

In conclusion, with millions of T2DM patients around the world, it is clear that more treatment options need to be made available to meet the demand of these patients. The field is currently pursuing DPP-4 inhibition to treat T2DM since it was identified as a good target that minimizes patient risk of low blood sugar levels. Though several DPP-4 inhibitors are on the market, improvements on these systems can be undertaken to improve patient quality of life.

To make progress toward this feat, computational and chemical intuition-driven methods were carried out using anagliptin, a currently available DPP-4 inhibitor, as a control and source of inspiration. Two different sites on anagliptin were modified during computer and hypothetical modeling—both of which were successful in the sense that they improved the docking score of anagliptin mean while meeting ADMET criteria.

Though the results of these studies are promising in developing new DPP-4 inhibitory pharmaceuticals, further testing of these molecules is required before their widespread use in drug formulations. These software tools generated information on drug candidates in a quick and efficient manner, but they have limitations in the sense that they cannot provide information about how the molecule behaves in the biological setting. Despite having great docking scores and ADMET properties based on our data, a drug might undergo transformations in the body that alter its efficacy and tolerability. Nonetheless, these tools were helpful in finding new candidates for the treatment of diabetes.

## AUTHOR INFORMATION

### Author Contributions

Research was designed by all authors; all experiments were carried out by L.C.R. The manuscript was written through contributions of all authors. All authors have given approval to the final version of the manuscript.

## ACKNOWLEDGMENT

Research reported in this publication was supported by UC Davis, the National Science Foundation Award Numbers 1827246, 1805510, 1627539, the National Institute of Environmental Health Sciences of the National Institutes of Health (NIH) under Award Number P42ES004699, UC Davis, NIH Award Number R01 GM 076324-11 and the Rosetta Commons. The content is solely the responsibility of the authors and does not necessarily represent the official views of the National Institutes of Health or National Science Foundation. This study was derived from a course based undergraduate research study conducted in Chemistry 130B at UC Davis.

## REFERENCES

1. National Diabetes Statistics Report, 2017; U.S. Dept of Health and Human Services. Centers for Disease Control and Prevention: Atlanta, GA, 2017.

2. Deaths: Final data for 2017. National Vital Statistics Reports; vol 68 no 9. National Center for Health Statistics: Hyattsville, MD: 2019.

3. Guasch, L.; Ojeda, M. J.; González-Abuín, N.; Sala, E., Cereto-Massagué, A.; Mulero, M.; Valls, C.; Pinent, M.; Ardévol, A.; Garcia-Vallvé, S.; Pujadas, G. PloS one. 2012, 7(9), 44971.

4. Jiang, J.; Ghosh, S. PDB101: Global Health: Diabetes Mellitus: Drugs: DPP4 inhibitors. https://pdb101.rcsb.org/global-health/diabetes-mellitus/drugs/dpp4-inhibitor/dpp4 (accessed Apr 13, 2019).

5. FDA Drug Safety and Availability. https://www.fda.gov/drugs/drug-safety-and-availability/fda-drug-safety-communication-fda-adds-warnings-about-heart-failure-risk-labels-type-2-diabetes (accessed Aug 29, 2019).

6. Raschi, E.; Piccinni, C.; Poluzzi, E.; Marchesini, G.; De Ponti, F. Acta Diabetologica. 2013, 50, 4, 569–577.

7. Elashoff, M.; Matveyenko, A. V.; Gier, B.; Elashoff, R.; Butler, P. C. Gastroenterology. 2011, 141(1), 150–156.

8. Piyush, G.; Kumar, J. S.Indian J of Pharmaceutical Education and Research. 2017, 51(4), 664–671.

9. Kalhotra, P.; Chittepu, V. C. S. R.; Osorio-Revilla, G.; Gallargo-Velázquez, T. Molecule. 2018, 23(6), 1368.

10. Berman, H. M.; Westbrook, J.; Feng, Z; Gilliland, G.; Bhat, T. N.; Weissig, H.; Shindyalov, I. N.; Bourne, P. E. Nucleic Acids Research. 2000, 28(1), 235–242.

11. RCSB PDB-3WQH: Crystal Structure of human DPP-IV in complex with Anagliptin. https://www.rcsb.org/structure/3WQH (accessed Apr 13, 2019).

12. RCSB PDB-3G0B: Crystal Structure of dipeptidyl peptidase IV in complex with TAK-322. https://www.rcsb.org/structure/3G0B (accessed Apr 13, 2019).

13. RCSB PDB-2RGU: Crystal structure of complex of human DPP4 and inhibitor. https://www.rcsb.org/structure/2RGU (accessed Apr 13, 2019).

14. RCSB PDB-1X70: HUMAN DIPEPTIDYL PEPTIDASE IV IN COMPLEX WITH A BETA AMINO ACID INHIBITOR. https://www.rcsb.org/structure/1X70 (accessed Apr 13, 2019).

15. The PyMOL Molecular Graphics System; Schrödinger, LLC: 2000.

16. FILTER; OpenEye Scientific Software: Santa Fe, NM.

17. BROOD, 3.1.1.2; OpenEye Scientific Software: Santa Fe, NM.

18. Gaussian 03; Gaussian, Inc.: Wallingford, CT, 2009.

19. Hawkins, P. C. D.; Skillman, A. G.; Warren, G. L.; Ellingson, B. A.; Stahl, M. T. J. Chem Inf. Model. 2010, 50, 4, 572–584.

20. McGann, M. J. Chem. Inf. Model. 2011, 51, 3, 578–596.

21. Chan, L. L.; Lidstone, E. A.; Finch, K. E.; Heeres, J. T.; Hergerother, P. J.; Cunningham, B. T. JALA Charlottesv Va. 2009, 14(6), 348–359.

22. Jiang, S.; Zhang, L.; Cui, D.; Yao, Z.; Gao, B.; Lin, J.; Wei, D. Scientific Reports. 2016, 6, 34750.

